# Integrative Genome-Wide Association Analysis Reveals Polygenic Basis and Novel Candidate Genes for Uncapping Behavior in Varroa-Resistant in Worker Bees

**DOI:** 10.64898/2025.12.05.692650

**Authors:** Peymaneh Davoodi, Mohammad Razmkabir

**Author notes:** Corresponding Authors: Peymaneh Davoodi, and, Mohammad Razmkabir.

## Abstract

Uncapping behavior of Varroa-infested brood cells in honeybees represents a key social immunity trait and a practical marker for breeding Varroa-resistant colonies. We applied genome-wide association analyses using both linear and logistic MLM-LOCO models (Leave one chromosome out) on Affymetrix 44K SNP data to dissect the genetic architecture of this behavior. Seven loci were identified in linear analyses, with partially overlapping signals in binary case-control models. Three SNPs consistently detected across both approaches highlight robust genomic markers for hygienic behavior. Candidate genes implicated in transcriptional regulation, neurodevelopment, cuticular integrity, and olfactory signaling suggest a coordinated molecular basis for behavioral resistance. These results underscore the polygenic nature of uncapping behavior and demonstrate how integrative GWAS frameworks can reveal complex traits in social insects, providing a foundation for marker-assisted selection to enhance colony resilience. Beyond apiculture, these results advance understanding of the genetic basis of cooperative defense traits in social animals.

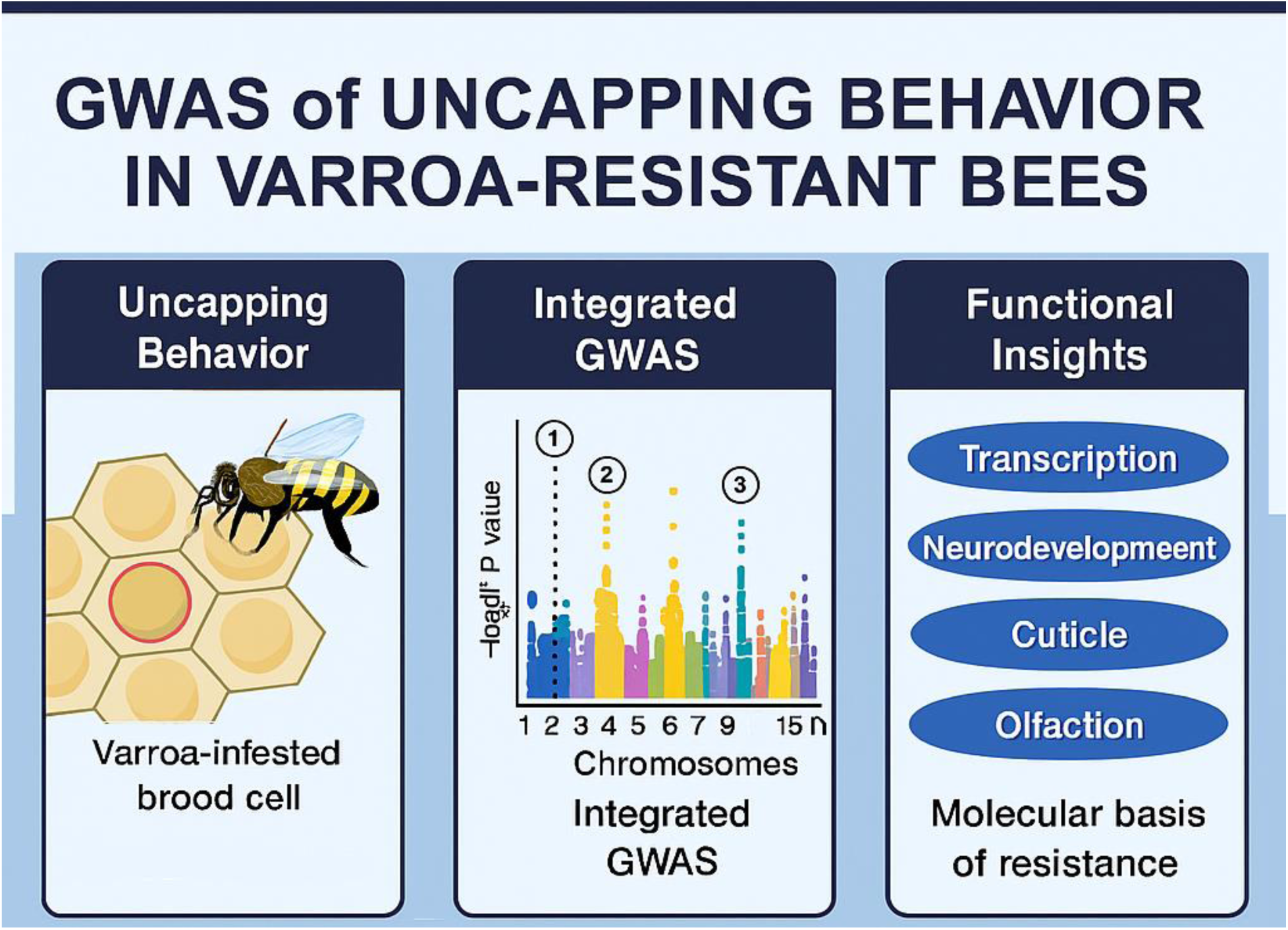

## Introduction

Honey bees (*Apis mellifera*) are among the most important pollinators in agricultural and natural ecosystems, contributing substantially to global food security and biodiversity. Yet, in recent decades, honey bee populations have experienced alarming declines, with colony losses reported worldwide. Multiple stressors contribute to these declines, among these, infestations by the ectoparasitic mite *Varroa destructor* remain one of the most significant threats to colony survival (Behrens et al. 2011; Guichard et al. 2020; Frazier et al. 2024; O’Connell et al. 2025).

*Varroa destructor* reproduces within sealed brood cells, feeding on the hemolymph of developing pupae and adult bees. Beyond direct parasitism, the mite acts as an efficient vector for numerous pathogenic viruses, including deformed wing virus, thereby amplifying colony morbidity and mortality. The global spread of *Varroa* has rendered most unmanaged colonies unsustainable, underscoring the urgent need for effective, durable resistance strategies (Guichard *et al*. 2020; Frazier *et al*. 2024).

One promising avenue lies in the natural defense mechanisms expressed by worker bees, collectively termed hygienic behavior. This suite of behaviors includes the detection, uncapping, and removal of diseased or parasitized brood (Spötter et al. 2016; Khan & Ghramh 2021). In the context of *Varroa* infestation, uncapping behavior is particularly critical: by opening mite-infested cells, workers disrupt mite reproduction and reduce infestation pressure in subsequent generations. This trait, often referred to as Varroa-sensitive hygiene (VSH), has been shown to vary among colonies and is heritable, making it a practical target for selective breeding (Mondet et al. 2015; Spötter *et al*. 2016; Spivak & Danka 2021; Guichard et al. 2022; Morin & Giovenazzo 2023).

Uncapping behavior is increasingly viewed as a genetically influenced component of social immunity, contributing to colony health and resilience. Colonies with strong hygienic traits show reduced pathogen loads and improved survival, underscoring the ecological and evolutionary importance of these behaviors (Mondet et al. 2020; Spivak & Danka 2021; Haider et al. 2025). Despite its importance, the genetic architecture underlying uncapping behavior remains incompletely understood. Early quantitative genetic studies suggested moderate heritability (Boecking et al. 2000), while more recent genomic approaches have revealed a polygenic basis (Davoodi & Razmkabir 2025; Eynard et al. 2025). Genome-wide association studies (GWAS) are powerful tools for dissecting complex traits, and SNP arrays have successfully been applied in honey bees to identify loci associated with hygienic behavior and Varroa-specific defense (Pritchard 2016; Spötter *et al*. 2016; Guichard et al. 2021; Mustafa et al. 2024). These findings reinforce the potential of marker-assisted selection in apicultural breeding programs (Cheruiyot et al. 2018).

Beyond trait association tests, recent work has emphasized the complexity of behavioral genetics in social insects. Hygienic behavior likely follows an Omnigenic or Omnifactorial model, in which numerous loci of small effect, distributed across the genome, contribute to trait variation using fixed linear association tests (Davoodi & Razmkabir 2025). Such complexity underscores the need for integrative analytical frameworks that can capture both major and minor genetic effects. Advances in statistical genetics have facilitated this effort. Mixed linear models with leave-one-chromosome-out (MLM-LOCO) approaches reduce confounding from population structure and relatedness, improving the robustness of GWAS signals (Zhang et al. 2010; Jiang et al. 2019; Davoodi et al. 2022b). Applying both linear and logistic models allows complementary perspectives on trait architecture, enhancing confidence in identified associations.

This study explores the genetic basis of uncapping behavior in honey bee workers using GWAS across continuous and categorical phenotypes. Applying MLM-LOCO models with the Affymetrix 44K SNP array, the research aims to (i) define the polygenic architecture of the trait, (ii) identify robust SNP markers consistently detected across analyses, and (iii) highlight candidate genes involved in social immunity. The resulting markers and gene candidates provide insights into mechanisms of mite resistance and support the development of breeding strategies for more resilient honey bee colonies.

## Materials and Methods

### Experimental data

#### Data Resources and Breeding Design

All colonies were produced through a controlled cross-breeding program designed to generate variation in the Varroa-specific defense trait of uncapping mite-infested brood cells at the Institute for Bee Research, Hohen Neuendorf (Spötter et al. 2012; Spötter *et al*. 2016).

#### Sampling, Colony Maintenance, and Phenotyping

A total of 22,000 worker bees were individually labeled with numbered tags to enable identification and behavioral tracking. Sampled workers descended from 10 distinct, labeled queens (families). Colonies were maintained in observation hives with transparent glass walls, allowing direct visual inspection of the comb. Standard husbandry protocols were applied consistently across colonies, including regulated temperature, feeding, and colony management practices. Observation hives were continuously video-recorded using infrared cameras for seven consecutive days, enabling uninterrupted monitoring of all tagged individuals. The focal trait was the Varroa-specific defense behavior of uncapping infested brood cells. Phenotypes were retrospectively assigned to cases and controls based on recorded behavior, following predefined definitions and quality-control criteria. Sampling was conducted across two consecutive years: in year one (label A871), 91 bees were genotyped, and in year two (label A1173), 128 bees were genotyped (Spötter *et al*. 2012). The two definitions of uncapping behavior in worker bees were used: In the context of the linear analysis of uncapping variation, it refers to the number of uncapping brood cells from 0 to 4 in certain time of experiment. Conversely, the binary measure of uncapping variation categorizes the behavior just into two distinct groups. In this definition, each worker bee is classified simply as either exhibiting the behavior (“uncapping-positive”/ healthy) or not exhibiting it (“uncapping-negative”/disease) (Spötter *et al*. 2012; Spötter *et al*. 2016; Davoodi & Razmkabir 2025).

#### Quality control

Overall, 44,000 SNPs were considered for the GWAS construction. During the quality control process, a total of 29,841 variants and 219 bee samples (females) were analyzed (Spötter *et al*. 2012). Of these, 29 samples and 1,254 variants were excluded due to missing genotyping data, 1,408 variants were removed following the Hardy-Weinberg exact test, and 12,082 variants were excluded based on the minor allele frequency threshold. Ultimately, 15,097 variants and 190 individuals successfully passed the filters and quality control (QC) and were found to be polymorphic in our tested populations.

#### LD score

Linkage Disequilibrium score (LD score) quantifies the amount of genetic correlation between variants in a given genomic region. The LD score for a specific SNP is calculated by summing the squared correlation coefficients (r²) between that SNP and all other SNPs within a defined region of 1000 base window. LD Score of 0 represents No detectable linkage disequilibrium, meaning the variant is independent and not correlated with nearby SNPs. A higher LD score for a SNP indicates that it is in LD with other SNPs, meaning it’s likely tagging a larger number of genetic variants (Yang et al. 2011).

#### Genome-Wide Association Studies

Genome-wide association studies (GWASs) on linear and binomial definition of uncapping trait in worker bees were performed using the GCTA software, applying the GCTA-LOCO (Leave-One-Chromosome-Out) linear mixed model (Yang *et al*. 2011). This approach uses the standard linear mixed model by including the additive effect of each SNP individually. Specifically, the following model was used for GWAS:

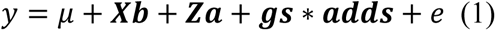

where *𝐲* is uncapping phenotype, *𝛍* is overall mean, *𝐗 𝐛* is matrix and its coefficients of fixed effects (queen, year and PC effects), *𝐙 𝐚* is matrix and its coefficient of random effects of polygenic effects derived from GRM, 𝒈 **s** is the genotype vector for **s**^th^ SNP, and 𝒂 𝒅 𝒅 **s** is the additive effect of **s**^th^ SNP and e is residual of the models. The genomic relationship matrix (GRM) is computed by excluding SNPs located on the same chromosome as the candidate SNP (LOCO), reducing confounding effects caused by local linkage disequilibrium (LD).

Significance thresholds for SNPs were determined using relaxed threshold of p-value 0.001 to identify potentially relevant effects. Manhattan plots were generated using the qqman package in R (v2024.09.0). The genomic data were subsequently analyzed in PLINK to assess LD pattern (Purcell et al. 2007). To define candidate genomic regions surrounding the significant genomic signals in the GWAS, a 200k bp window upstream and downstream of the significant SNPs were searched (Donthu et al. 2024).

#### Comparative Genomics Analysis for Gene Annotation

A synteny comparison between two assemblies of the Western honey bee (Apis mellifera), including Amel_4.0 (GCF_000002195.2) and Amel_HAv3.1 (GCF_003254395.2) was illustrated in Fig 6. Each assembly is depicted as a linear sequence of 16 chromosomes (labeled LG01–LG16). The green bands represent aligned regions or “syntenic blocks” where the genome structure between assemblies is highly similar, while blue lines indicate rearrangements and misassemble artifacts. In comparative view, synteny plot is essential for comparing assemblies (Fig 6) revealing and correcting assembly errors, and supporting downstream genomic annotation efforts (Polevikov & Kolmogorov 2019).

#### Proportion of Variance Explained

The Proportion of Variance Explained (PVE) calculation was utilized to estimate the contribution of significant SNPs identified through GWAS to the phenotypic variance of the uncapping trait in worker bees. This was determined using the formula established by Shim et al. (2015) (Shim et al. 2015; Han et al. 2024).

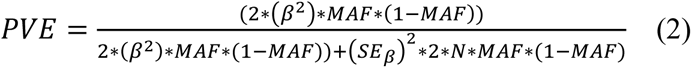

where 𝛽 represents the estimated effect size of the SNP, MAF is the minor allele frequency, 𝑆 𝐸 _𝛽_ is the standard error of the effect size, and N is the sample size. This formula accounts for both the genetic effect and sampling variance, providing an accurate estimation of the variance explained by each SNP.

#### Integrative Gene Ontology Enrichment and Network Analysis

Initially, previously published studies (2005-2025) related to Varroa mite resistance and uncapping behavior were surveyed using a module of pathway/process/disease in the STRING database (Table 1) (Davoodi & Ehsani 2019; Szklarczyk et al. 2021; Szklarczyk et al. 2023). Then genes identified in each study were extracted and subsequently merged with candidate genes obtained from the present study. The resulting gene set was curated and utilized as an integrated dataset for network analysis and gene ontology enrichment through the STRING database. Gene Set Enrichment Analysis (GSEA) was subsequently conducted using Honeybee STRING 12.0 to identify biological pathways significantly enriched with the SNPs of interest, and FDR cutoff of 0.01 was applied (Szklarczyk *et al*. 2021; Szklarczyk *et al*. 2023).

**Table 1.** The list of published articles from STRING with the number of introduced genes associate<u>d with varroa resistance</u>.

| PMID | Gene | Year | References |
| --- | --- | --- | --- |
| 39560851 | 2 | 2024 | (Kabakcı et al. 2024) |
| 36725182 | 5 | 2023 | (Zheng et al. 2023) |
| 34729804 | 3 | 2022 | (Guichard *et al.* 2022) |
| 33454582 | 2 | 2021 | (Flores et al. 2021) |
| 35087998 | 25 | 2020 | (Kaskinova et al. 2020) |
| 32537655 | 6 | 2020 | (Noël et al. 2020) |
| 29728057 | 8 | 2018 | (Mondet et al. 2018) |
| 28740210 | 15 | 2017 | (Annoscia et al. 2017) |
| 28081188 | 7 | 2017 | (Koleoglu et al. 2017) |
| 28154949 | 7 | 2017 | (Hamiduzzaman et al. 2017) |
| 29088251 | 7 | 2017 | (Tesovnik et al. 2017) |
| 26000641 | 18 | 2015 | (Mondet *et al.* 2015) |
| 25366780 | 5 | 2014 | (Ji et al. 2014) |
| 26466617 | 7 | 2012 | (Aronstein et al. 2012) |
| 23133626 | 4 | 2012 | (Tsuruda et al. 2012) |
| 22497859 | 4 | 2012 | (Gregorc et al. 2012) |
| 23021491 | 3 | 2012 | (Parker et al. 2012) |
| 23133594 | 2 | 2012 | (Arechavaleta-Velasco et al. 2012) |
| 18578863 | 8 | 2008 | (Navajas et al. 2008) |
| 16299597 | 3 | 2005 | (Gregory et al. 2005) |

## Results

Using the Affymetrix 44K SNP array and LOCO mixed linear models, we controlled for population structure and improved resolution compared with earlier fixed-model analyses. This approach revealed consistent association signals across both continuous and categorical phenotypes, enabling the identification of robust SNP markers and candidate genes linked to uncapping behavior.

### IBS Matrix Heatmap Analysis Before and After QC

The IBS heatmaps (Fig. 1) illustrate how quality control improved the resolution of population genetic structure. Raw data showed scattered high-similarity signals, likely due to relatedness, stratification, or technical artifacts, resulting in noisy patterns without clear clustering. After removing outliers and low-quality samples, the IBS matrix revealed distinct block-diagonal structures and coherent dendrograms, indicating genetically homogeneous clusters. These results emphasize the importance of rigorous filtering for accurately delineating population substructure in low-quality datasets.

**Fig 1.**
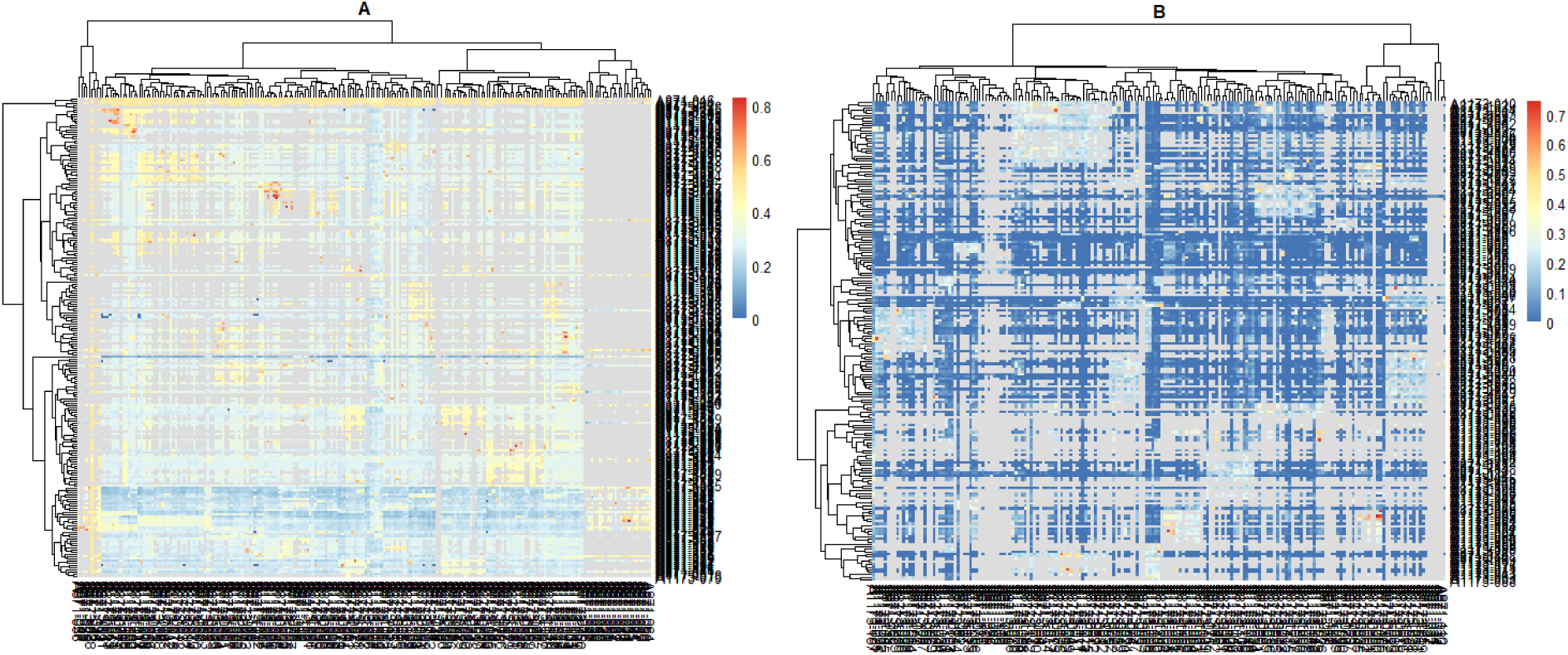
Heatmap of squared format of the IBS Matrix. **left**: raw data and **right**: after outlier deletion

### Allele frequency and LD Score

The histogram of allele frequency shows a uniform scatter in the frequencies from 0 to 0.5 in the studied SNPs after quality control. A local peak at a frequency of around 0.1 is also observed among the studied SNPs (Fig 2, top). The histogram of Linkage Disequilibrium scores (LD score) is shown in Fig 2, bottom. In the present study, the maximum value is 10, and the peak is seen at 2. LD score analysis revealed generally low linkage disequilibrium, with each SNP tagging fewer than 10 variants, limiting GWAS resolution. Common variants are easier to detect but typically have modest effects, while rare alleles may exert stronger impacts yet require larger cohorts or specialized methods. These findings highlight the importance of considering both LD structure and allele frequency when interpreting association signals and designing future studies (Panagiotou et al. 2010; Visscher et al. 2017; Uffelmann et al. 2021). In addition, LD scores are crucial in GWAS to account for polygenic effects and improve statistical power (Christoforou et al. 2012). The maximum value of 10, and the peak at 2 (Fig 2. bottom) indicates the condition of very diverse populations like honeybees.

**Fig 2.**
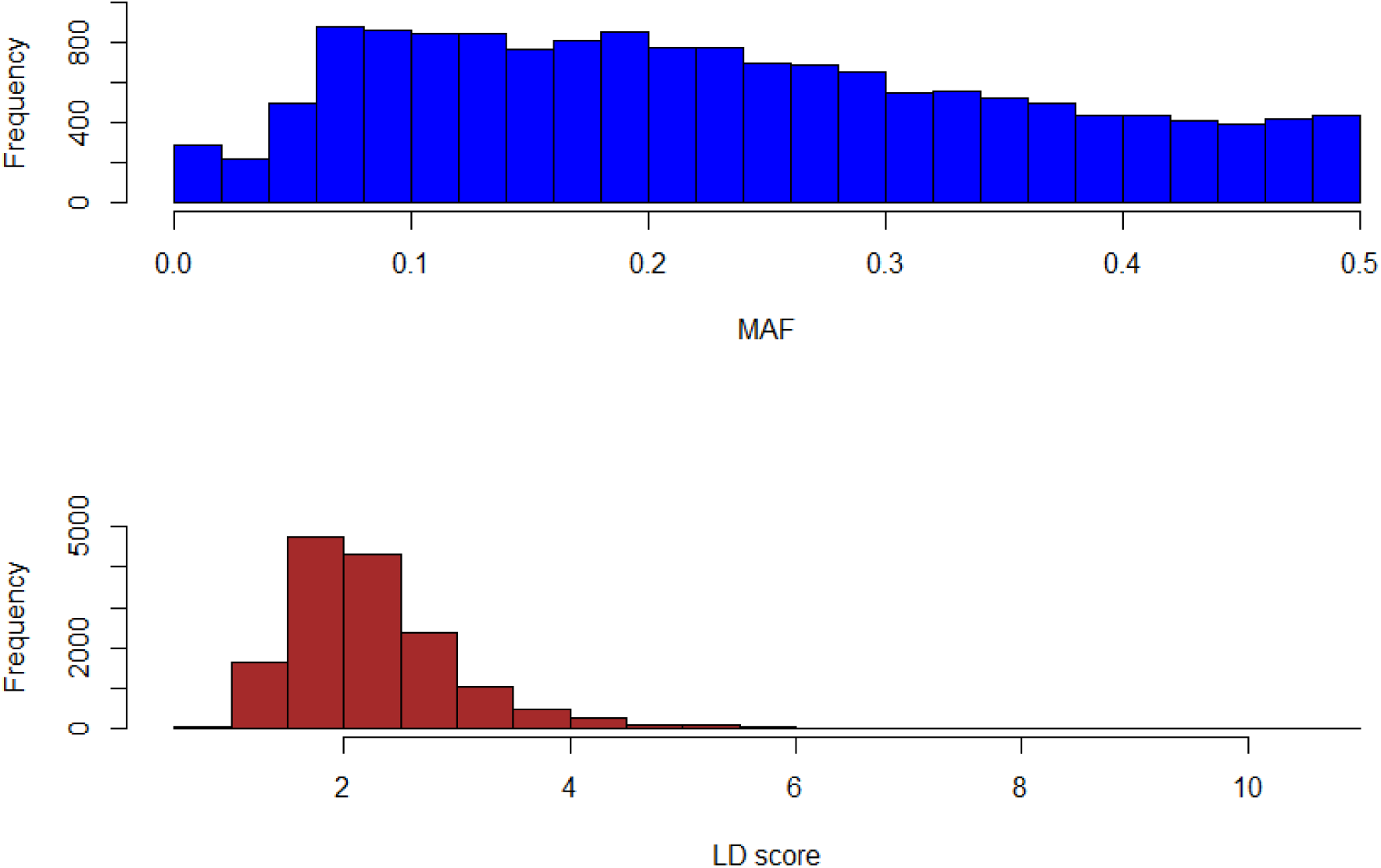
top: Allele frequency, bottom: LDscore frequency in data

### Principal Component Analysis (PCA)

Principal Component Analysis (PCA) (Fig 3.) were applied for managing population structure. It identifies and accounts for genetic variations due to ancestry, which can otherwise lead to false positives or inflated signals in GWAS if not considered (Davoodi *et al*. 2022b; Grinde et al. 2024). Including first principal components explaining 8.80 of variation as covariates in the GWAS model, can control for population stratification and obtain more accurate results.

**Fig 3.**
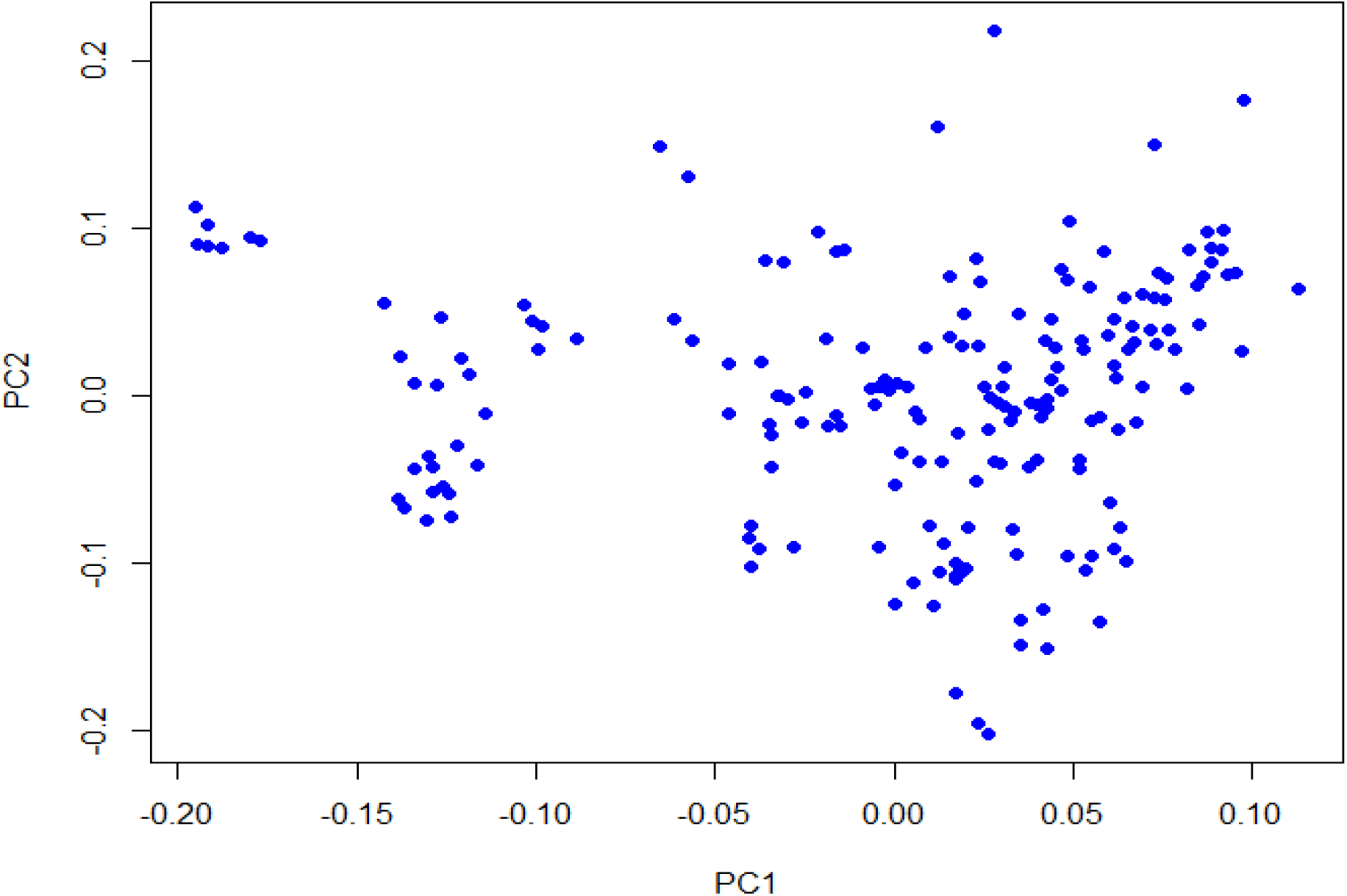
The PC plot showing a slight clustering with 8.8% explained variance

### Quality Control of Results

The Quantile-Quantile plot (QQ plot) in GWAS is used to assess whether the distribution of observed p-values aligns with the expected null distribution. It helps detect inflation or bias in association results which approves good power of test in both linear and case-control definition of uncapping behavior in worker bees (Fig 4).

**Fig 4.**
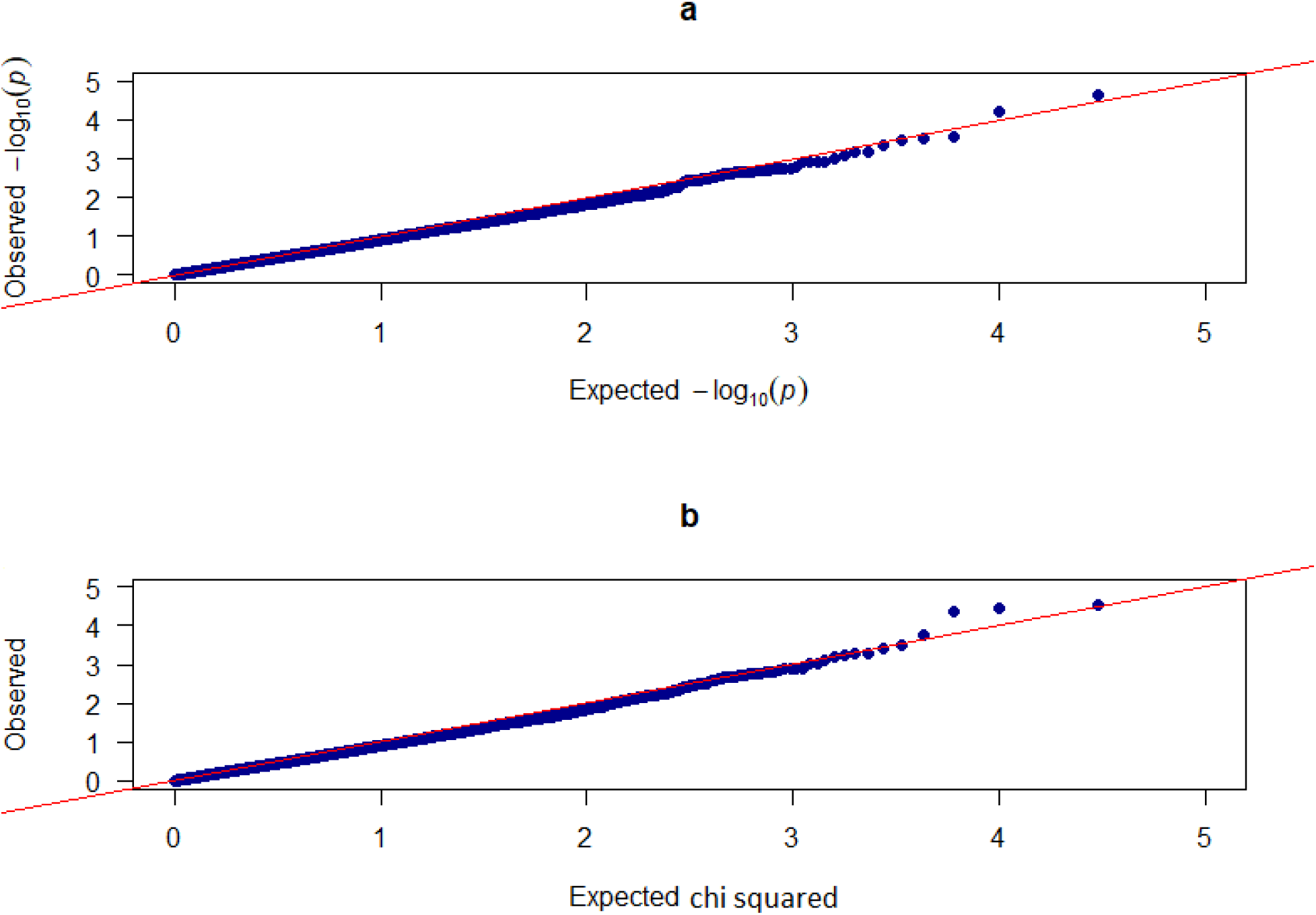
QQ plot of p-values extracted from GWASs, a) for Linear GWAS, b) For logistic GWAS. Diagonal red line (Expected Distribution)

### Two Scenarios of Genome-Wide Association Study

In linear GWAS, 7 SNP markers of *AMB-00377381*, *AMB-00333896*, *AMB-00037196*, *AMB-00230239*, *AMB-00147227*, *AMB-00732866*, and *AMB-00017454*, located in 4, 14, 12, 7, 2, 5, and 10 chromosomes, respectively, are detected to be associated with uncapping behavior (Fig 5a). On the other hand, in the logistic GWAS, 7 SNP markers of *AMB-00333896*, *AMB-00377381*, *AMB-00037196*, *AMB-00491306*, *AMB-00893206*, *AMB-00901806*, and *AMB-00794542* were positioned on chromosomes 14, 4, 12, 13, 6, 8, and 1, which are defined as top-associated SNPs with uncapping behavior in this study respectively (Fig 5b).

**Fig 5.**
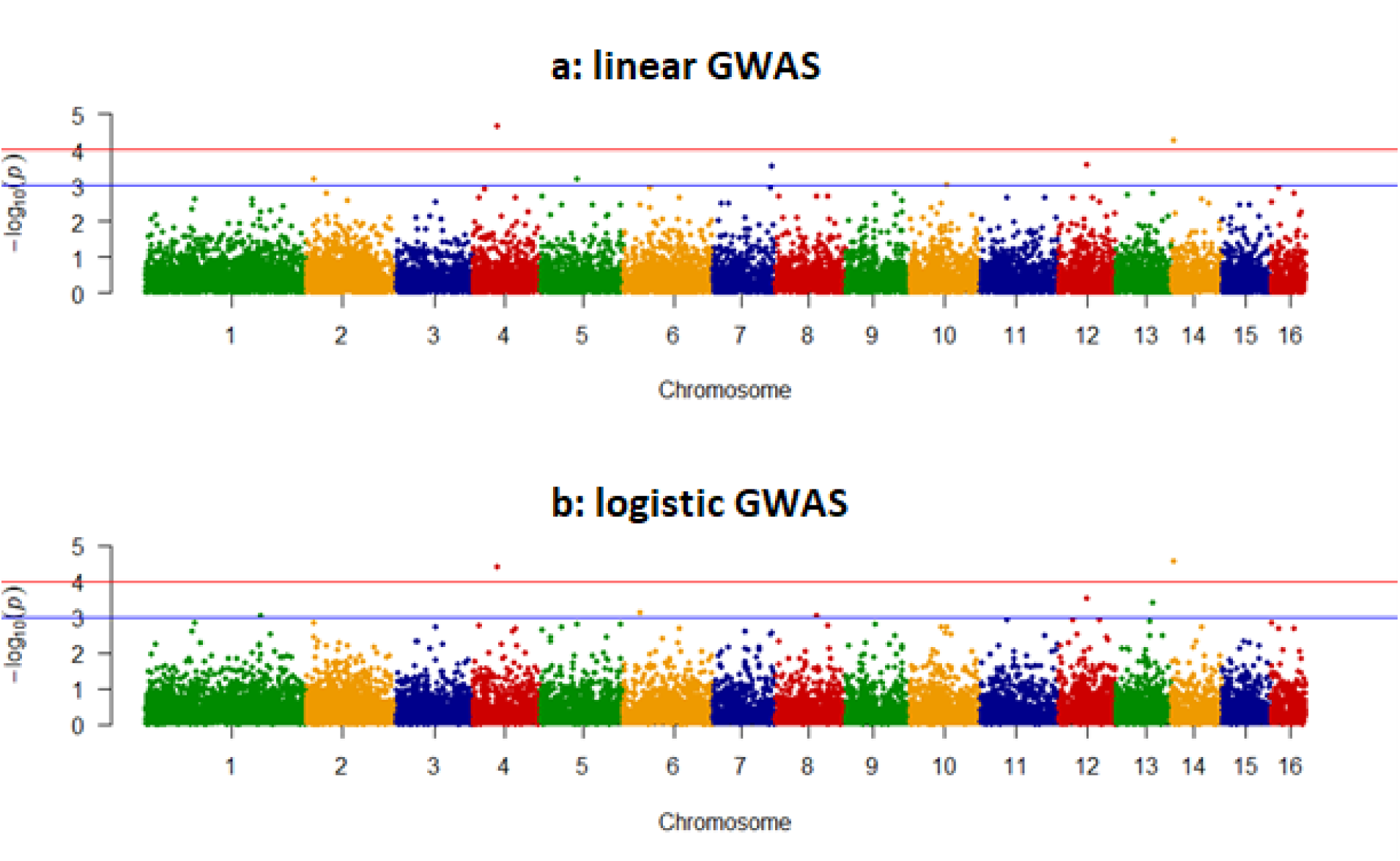
Manhattan plots to visualize the results of testing about 15,000 of genetic variants (SNPs) for association with a trait of uncapping brood cells. X-axis: Represents the physical position of SNPs along the chromosomes. The chromosomes are usually arranged sequentially. Y-axis: Represents the negative logarithm (base 10) of the p-value or chi-square value for each SNP’s association with the trait of interest. Significance Threshold: Horizontal lines indicate a significance and suggestive threshold. SNPs that fall above this line are considered statistically associated.

**Fig 6.**
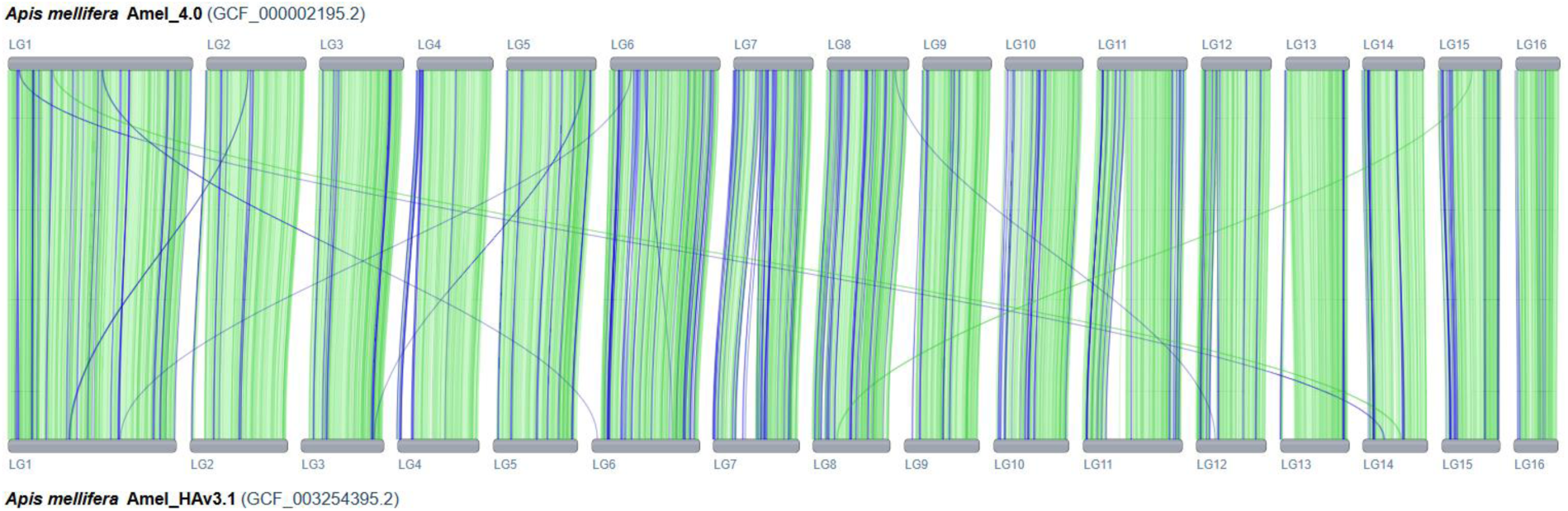
Comparative Genomics plot: Synteny between Apis mellifera two assemblies of Amel_0.4 and Amel_HAv3.1. Chromosomes are labeled as LG1 to LG16, green bands represent syntenic blocks, blue lines indicate misassemble artifacts.

Table 2 presents the findings of a genetic association analysis. Each row corresponds to a unique statistical model used to evaluate the association of a specific single-nucleotide polymorphism (SNP) with the trait of interest. The Gene column identifies the nearby gene(s) associated with each SNP, providing insights into potential biological significance. Among annotated genes with significant SNPs, five genes of *LOC100576831*, *LOC113219138*, *LOC551667*, *LOC100577335*, *LOC102655281*, and *LOC725477* were predicted protein-coding genes with unknown function. In conclusion, the functional annotation of genes harboring significant SNPs has uncovered a diverse array of molecular players potentially contributing to cellular regulation, development, immunity, and metabolism. Among these, several protein-coding genes with previously unknown roles, *LOC100576831*, *LOC113219138*, *LOC551667*, *LOC100577335*, *LOC102655281*, and *LOC725477*, highlight the genomic landscape yet to be fully explored.

### Overlapping Results between Two GWAS Scenarios

The Venn network at the SNP level (Fig 7) shows SNPs overlap between two GWAS scenarios: a Linear model design and a logistic model design, and the unique associated SNPs in each model design. Three shared SNPs including *AMB-00037196*, *AMB-00333896* and *AMB-00377381* were detected between two mentioned GWAS scenarios.

**Fig 7.**
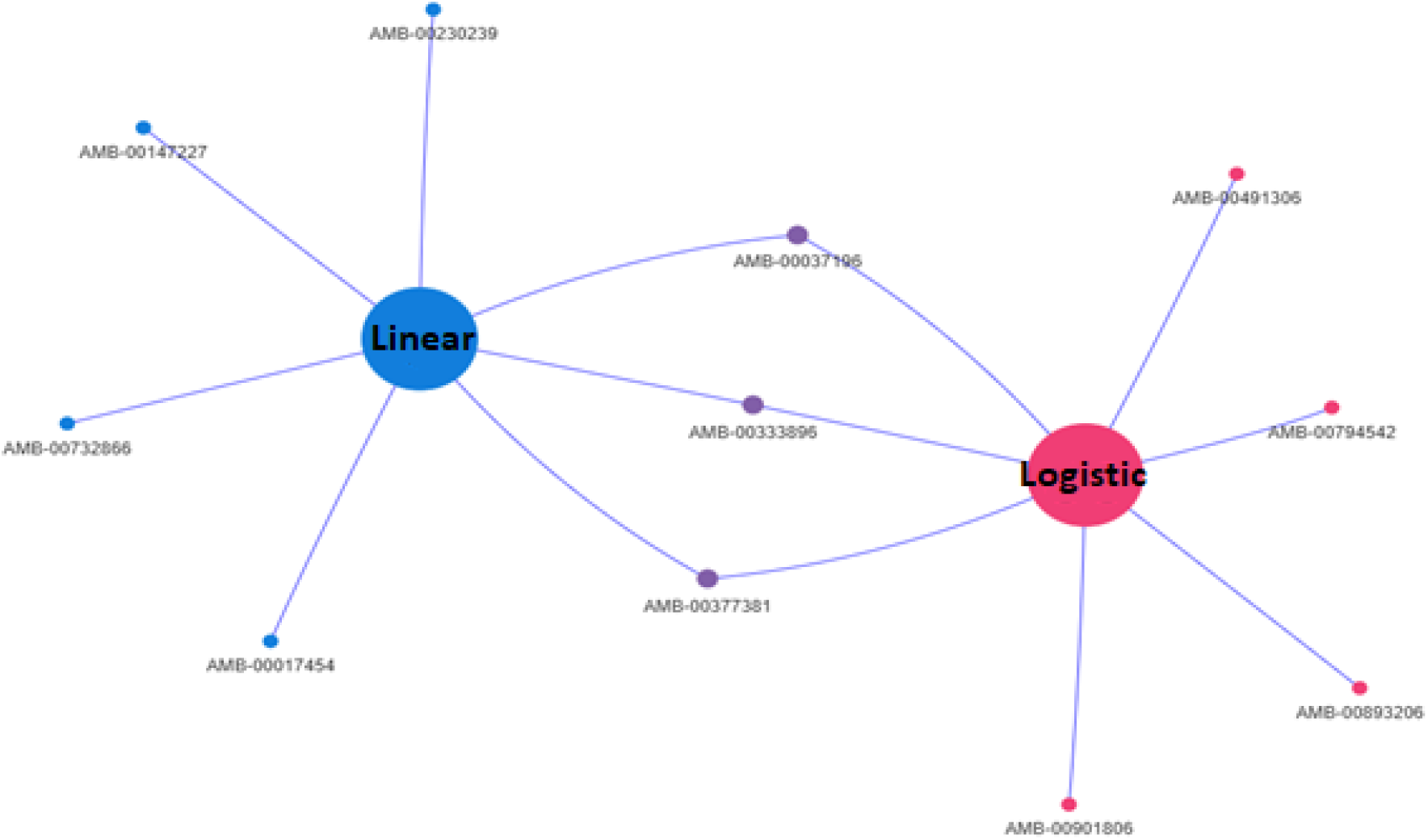
Overlapping SNPs between to GWAS scenarios of linear and logistic model

The Venn network at the associated gene level (Fig 8) shows Genes overlap between two GWAS scenarios of linear and logistic models, and the unique associated genes in each model design (17 and 24 genes). Nine overlapping genes including *LOC727092*, *LOC100577335*, *LOC100577295*, *LOC408803*, *LOC102655281*, *timeout*, *Mir6054*, *Cpr97Ea*, and *LOC411023* were detected that are shared in both the linear and logistic GWASs.

**Fig 8.**
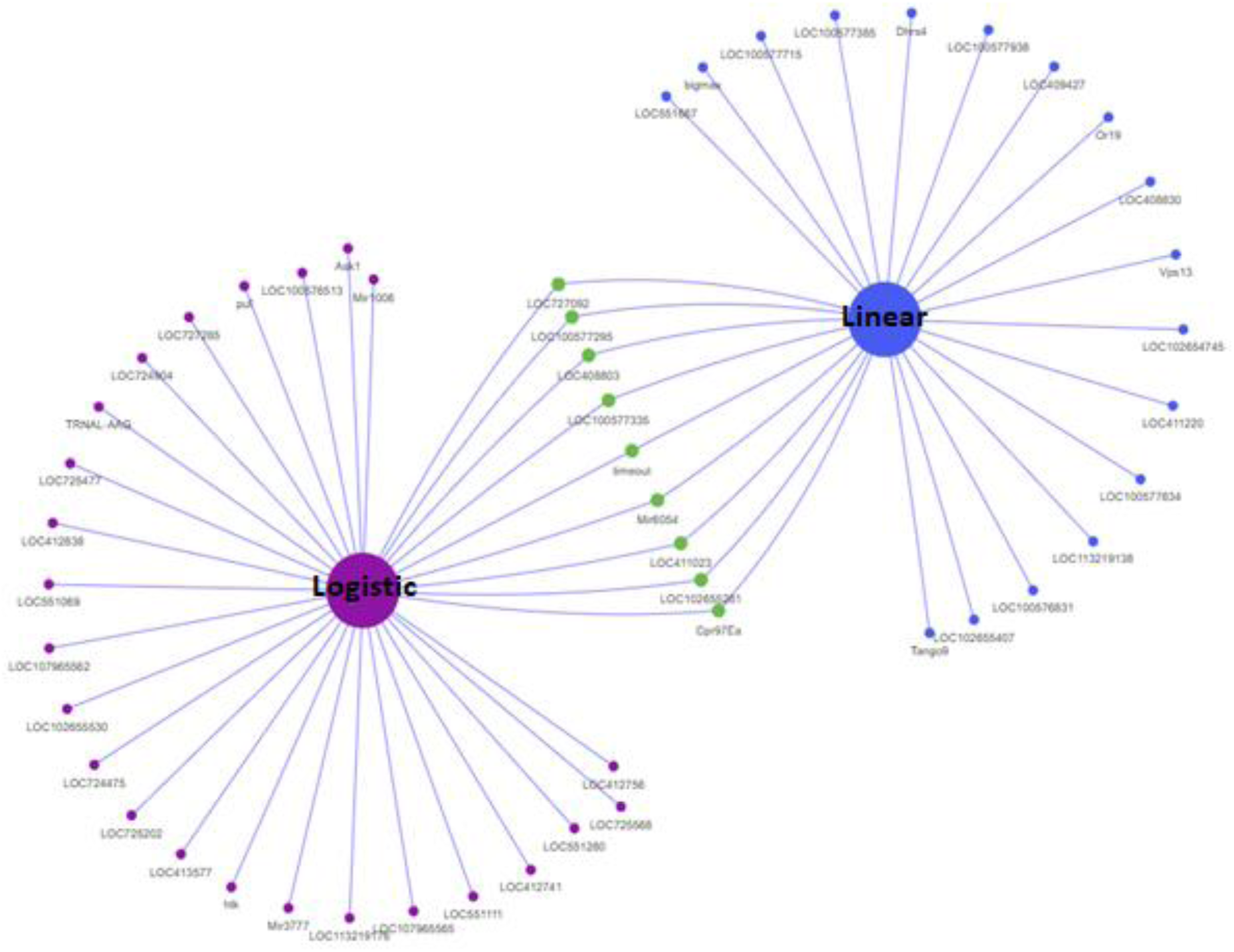
Overlapping genes between to GWAS scenarios of linear and logistic model

### Gene Ontology Enrichment and Integrative Network Analysis

To gain functional insight into genes associated with the uncapping trait and varroa resistance in honey bees, we constructed an integrated protein–protein interaction (PPI) network comprising 100 nodes and 537 edges. The network exhibited an average node degree of 10.7, indicating high connectivity among proteins. Additionally, the average local clustering coefficient was 0.482, suggesting that proteins tend to form functionally relevant clusters. The number of observed edges significantly exceeded the expected 170 for a random network of similar size, and the PPI enrichment p-value was < 1.0e^−16^, confirming that the observed interactions are biologically meaningful. These results underscore the functional coherence of the identified genes and support their potential involvement in molecular pathways linked to uncapping behavior (Fig 9).

**Fig 9.**
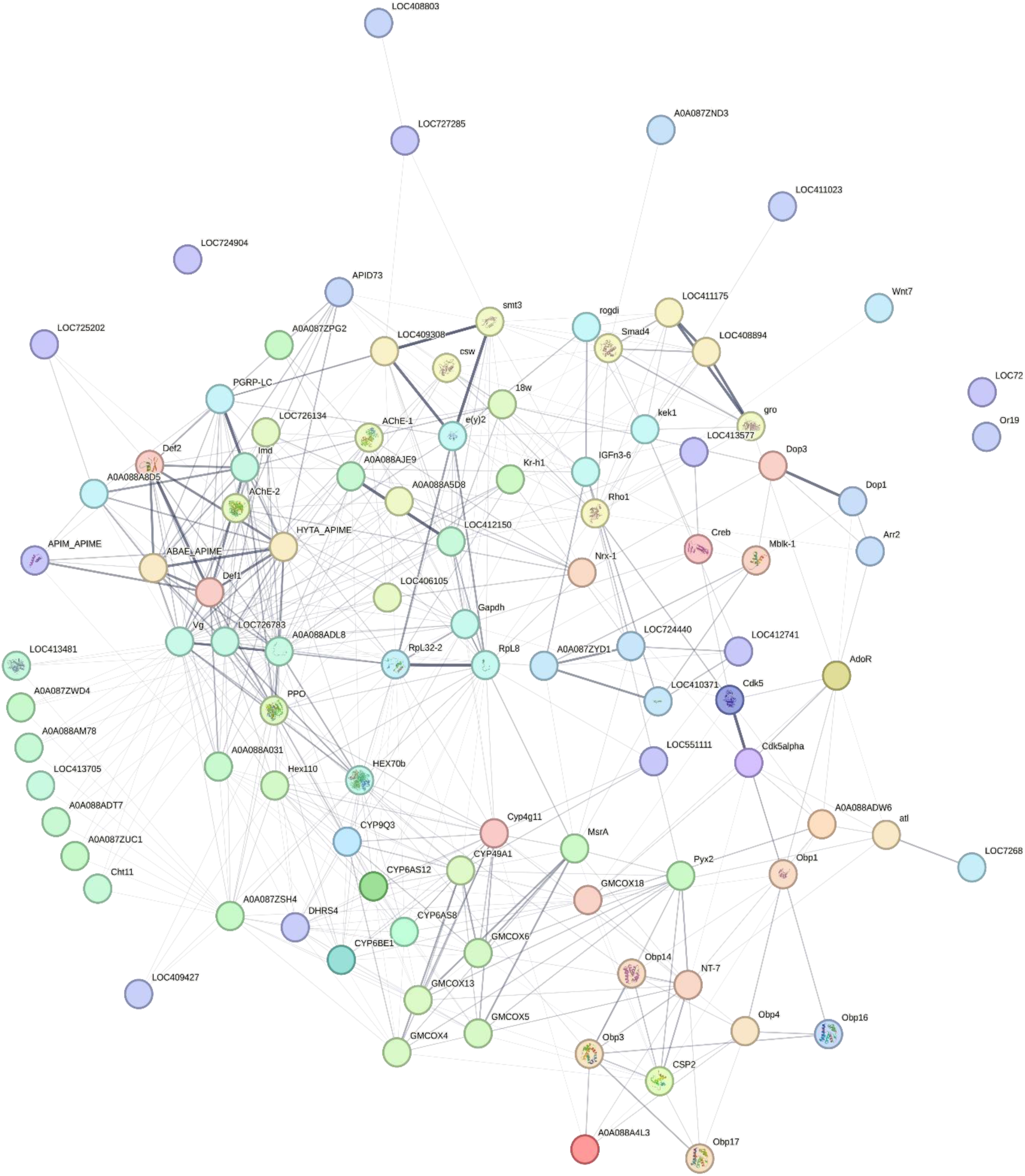
Integrative gene-gene network of genes associated with Varroa resistance

The KEGG pathway enrichment analysis revealed a multifactorial biological basis underlying Varroa resistance, characterized by the significant involvement of metabolic and signaling pathways (Fig 10a).

**Fig 10.**
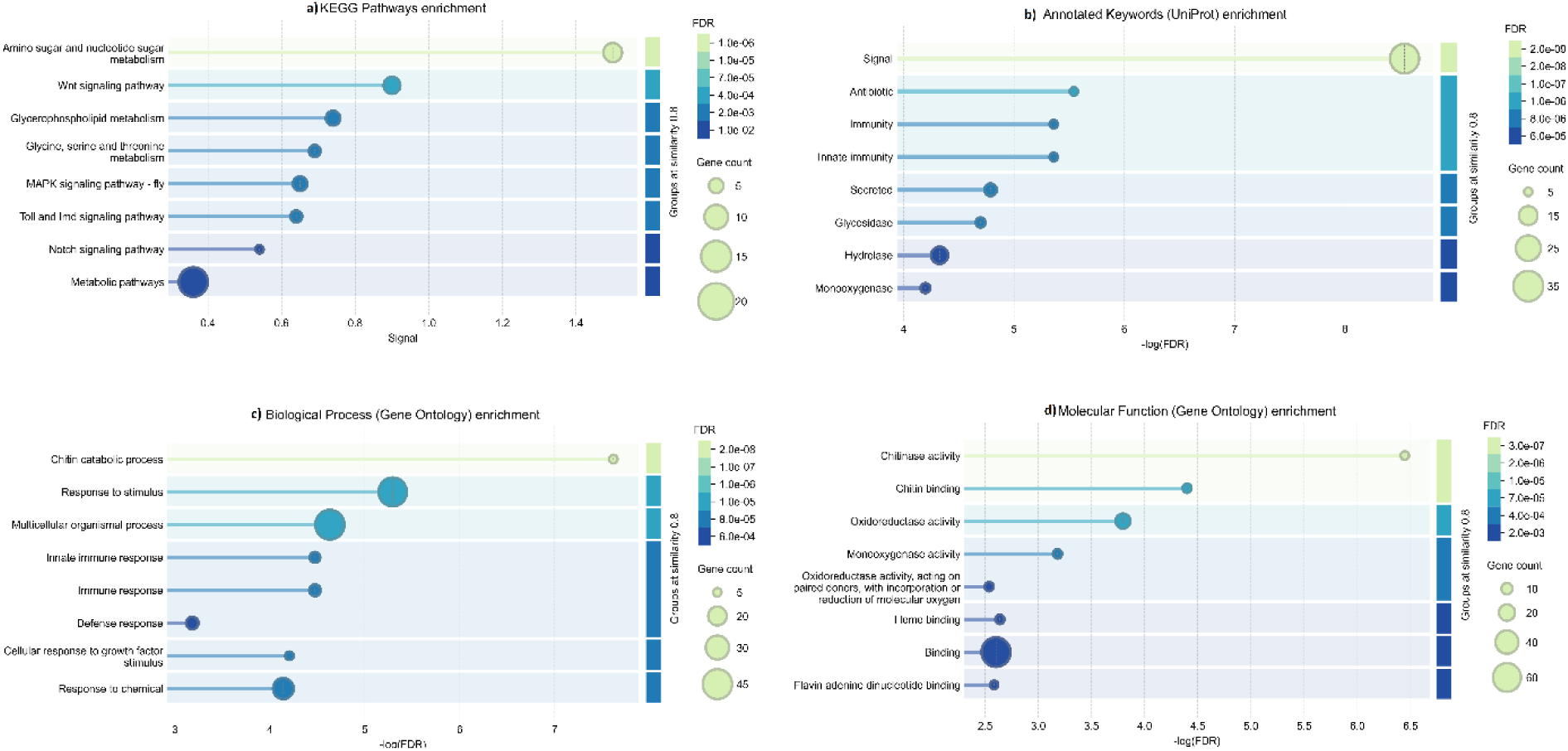
Integrative Gene Ontology (GO) enrichment of genes associated with Varroa resistance. a) KEGG pathway enrichment, b) Annotated keywords, c) Biological process (BP) and d) Molecular function (MF) enrichment

The annotated keyword enrichment analysis for Varroa resistance highlights a biologically diverse repertoire of genes contributing to host defense mechanisms (Fig 10b).

The Gene Ontology enrichment analysis revealed a distinct functional landscape of biological processes associated with Varroa resistance, marked by significant representation across immune, developmental, and stress-responsive pathways (Fig 10d).

Enrichment analysis revealed strong representation of metabolic pathways, including carbohydrate processing and biosynthesis, suggesting links to energy mobilization and structural integrity in resistant phenotypes. Key signaling cascades—Wnt, MAPK, Notch, and Toll/Imd—were also involved, indicating coordinated regulation of development, immunity, and stress responses. Notably, the Toll and Imd pathways play a central role in insect innate immunity, likely contributing to enhanced pathogen defense (Yang et al. 2023).

Further enrichment highlighted pathways supporting membrane remodeling and biochemical adaptation (glycerophospholipid and amino acid metabolism), alongside extensive signaling activity (33 genes) that may facilitate rapid immune and stress responses. Genes annotated with Immunity/Innate immunity, Antibiotic, and Secreted functions point to heightened innate defense and antimicrobial peptide secretion, while enrichment in hydrolase, glycosidase, and monooxygenase activity suggests roles in metabolic reprogramming, detoxification, and structural remodeling.

At the process level, the strongest enrichment was observed in multicellular organismal processes (44 genes), with additional representation in response to stimulus (42), response to chemical (24), and immune/innate immune responses (10 and 9), reflecting dynamic environmental sensing and defense activation. Processes such as defense response, cellular response to growth factors, and chitin catabolism further underscore adaptations in communication, tissue remodeling, and parasite interaction.

Enrichment analyses also revealed extensive binding functions (64 genes), alongside specialized activities such as chitinase/chitin binding, oxidative and detoxification pathways (oxidoreductase, monooxygenase, heme binding, FAD binding), all pointing to structural remodeling, enzymatic versatility, and redox resilience in Varroa resistance. These molecular functions suggest an integrated defense network at the cellular level.

## Discussion

Unlike our earlier work (Davoodi & Razmkabir 2025b), which relied on a fixed linear model implemented in PLINK, the present study employed a mixed linear model framework with explicit control for population structure. Using GCTA, we incorporated multiple principal components together with a genomic relationship matrix to account for relatedness among individuals (Yang *et al*. 2011).

The discovery of two microRNAs, *Mir6054* and *Mir1006* (Table 2), underscores the importance of post-transcriptional regulation in the genetic control of uncapping behavior and resistance to *Varroa destructor*. These microRNAs likely function as modulators within broader regulatory networks, contributing to the fine-tuning of social immunity pathways in honey bees. Their involvement highlights an additional layer of molecular complexity, suggesting that small RNAs may play a pivotal role in coordinating behavioral defenses against parasitic stress.

**Table 2.**
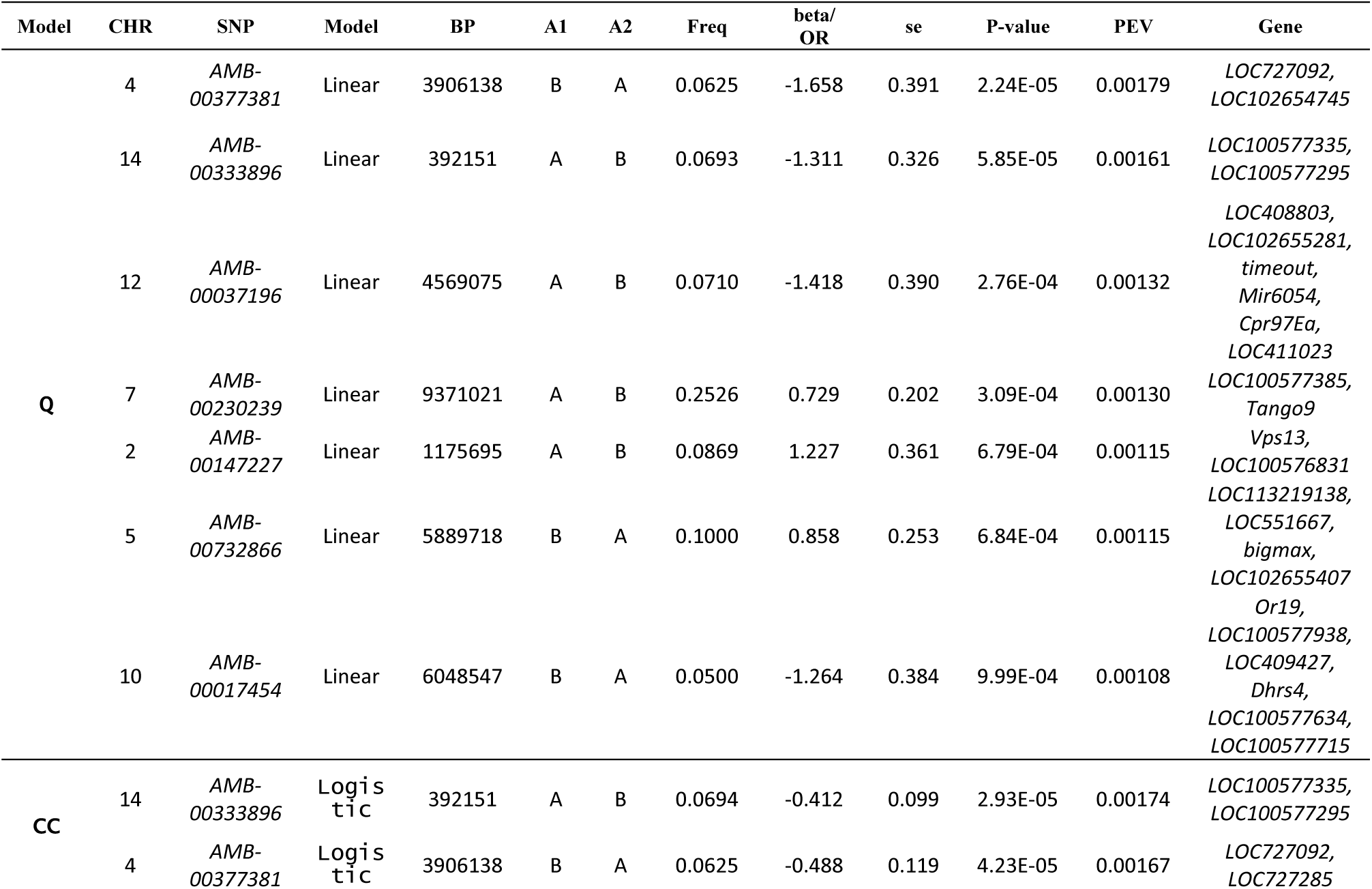
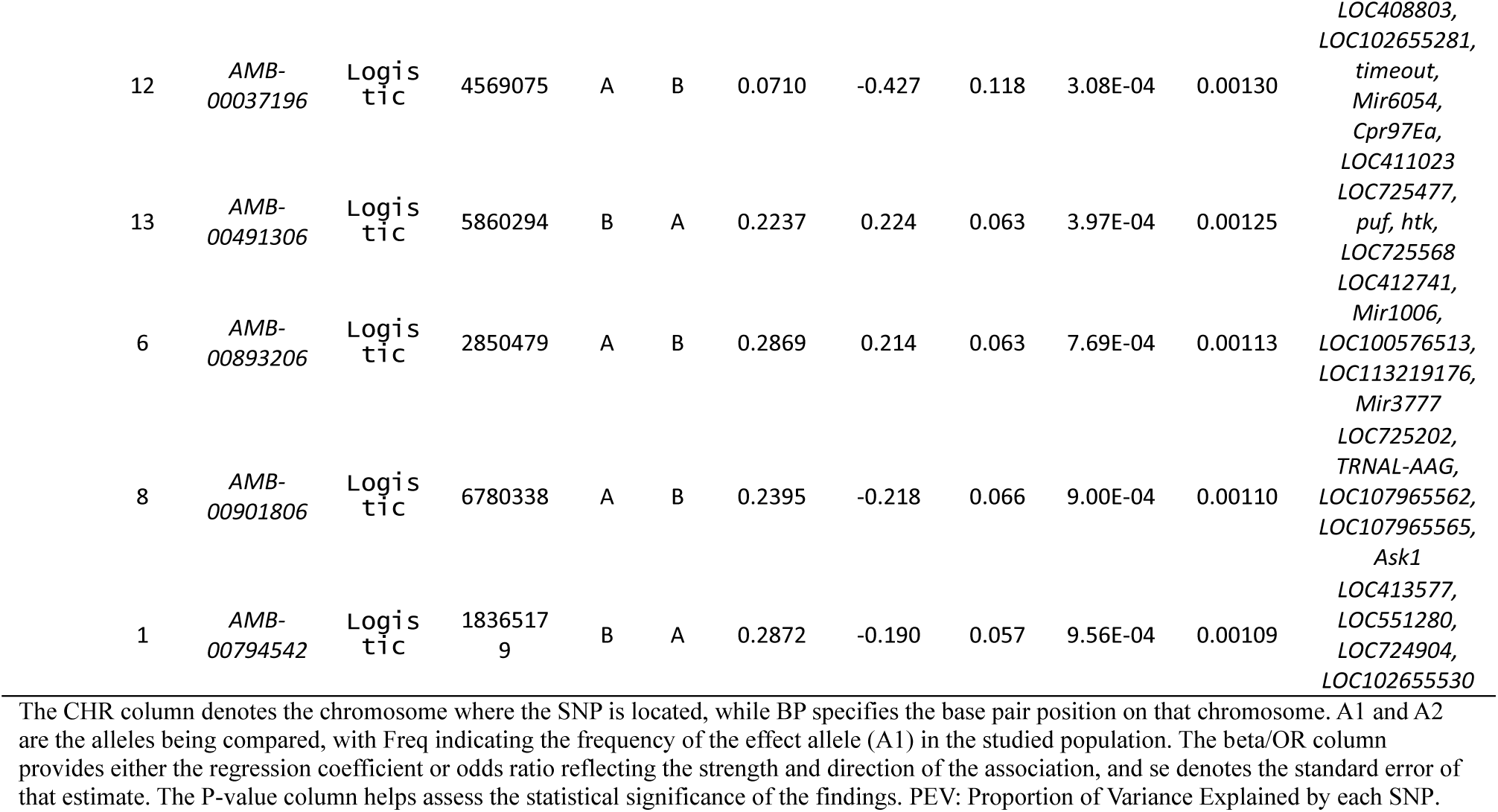
Summary results of genome-wide associations and gene annotation in two models with details.

Furthermore, *LOC727092*, a Nubbin-like transcription factor involved in developmental regulation via the POU domain, *LOC408803* may play a role in neural or sensory processes, and *Cpr97Ea* is a cuticular protein that has a structural role in the exoskeleton (Mallick & Eleftherianos 2024). In Drosophila, POU and homeodomain proteins serve a crucial immuno-protective function in barrier epithelia. Additionally, POU factor isoforms may function as molecular regulators, fine-tuning the expression levels of immune effectors. These factors also play a key role in managing host-microbe interactions within the gut (Tang & Engström 2019). It is reported that Drosophila nubbin (nub) is expressed in adult midgut progenitor cells, which promotes differentiation, functions as a tumor suppressor, and helps maintain intestinal stem cell proliferation (Tang et al. 2018). The gene *LOC411023* is also potentially linked to immune response. In addition, *Tango* appears to regulate vesicle trafficking within the Golgi organization, *Vps13*, a vacuolar protein sorting component, plays a role in intracellular transport, and *Bigmax* (MIX) is likely associated with growth regulation (Havula et al. 2013).

Furthermore, *Tango* family genes encode transmembrane proteins that facilitate the secretion of large lipoproteins, such as chylomicrons and very low-density lipoproteins (VLDLs), from the endoplasmic reticulum (Pfeffer 2016). According to the previous studies on insects, *Vps* proteins play a pivotal role in regulating vesicular trafficking, essential for maintaining cellular physiology, and are often exploited by various exogenous pathogens (Li & Blissard 2015; Vonk et al. 2017). *Or19* is an odorant receptor involved in sensory perception and olfaction. Studies have shown that odorant receptors are linked to behavioral traits in social honeybee and bumblebee species (Robertson & Wanner 2006; Gomez Ramirez et al. 2023).

*Dhrs4* functions as a dehydrogenase/reductase participating in metabolic processes specifically retinol metabolism (Hofmann et al. 2016). *Puf* is an RNA-binding protein that regulates mRNA stability and translation (Fischer & Olivas 2018). *Htk* is potentially a kinase-related protein that may contribute to signaling pathways (Scheiner et al. 2003). Additionally, *TRNAL-AAG* is a tRNA responsible for lysine translation (Avcilar-Kucukgoze & Kashina 2020). Lastly, *Ask1* is an apoptosis signal-regulating kinase that plays a critical role in stress response and cell death (Hattori et al. 2009).

LOC727092 encodes a nubbin-like transcription factor with essential roles in immune protection, epithelial integrity, and gut homeostasis in *Drosophila* via POU domain regulation. Alongside the developmental and neural relevance of LOC408803 and the structural contributions of Cpr97Ea, this highlights diverse gene functions. LOC411023, linked to immune response, together with Tango and Vps13, illustrates complex regulatory networks in vesicle trafficking and intracellular transport, critical for cellular physiology and pathogen defense. Additional genes, including odorant receptor Or19, metabolic enzyme Dhrs4, translational regulator Puf, kinase Htk, tRNA TRNAL-AAG, and apoptosis signal-regulating kinase Ask1, contribute to sensory perception, metabolic balance, mRNA stability, signal transduction, and stress responses. Collectively, this gene set reveals the interplay of developmental, immune, and regulatory processes, providing a foundation for functional validation and insights into their biological significance and evolutionary conservation in insects. (Ma et al. 2025).

Then, to better understand the genetic basis of uncapping behavior in honeybee workers, the contribution of individual SNPs to phenotypic variation using PEV estimates were quantified. Each PEV value represents (Table 2) the proportion of phenotypic variance in uncapping behavior that’s explained by an individual significant SNP (Zhu & Zhou 2020). These values are quite small, ranging from 0.11–0.18%, which is common in complex traits influenced by many genes and environmental factors. The cumulative genetic contribution of the identified SNPs was estimated to be approximately 1.5% by summing their PVE values. Collectively, though, these SNPs may still provide valuable insight into genetic architecture, especially if combined into polygenic scores or used for marker-assisted selection in the future.

Additionally, in GWAS research, identifying overlapping SNPs (Fig 7) and genes (Fig 8) helped to reveal genetic variants that are relevant regardless of whether a trait is analyzed linearly or dichotomously. Methods like polygenic overlap models can quantify the number and significance of such shared SNPs or genes, going beyond just genetic correlation between models and discovers more about genetic underlying of studied trait. The enrichment results collectively indicate that Varroa-resistant individuals rely on integrated signaling, enzymatic modulation, immune activation, and structural recalibration to counter parasitic threats. However, functional interpretation in honeybees is complicated by limited genome annotation, small GWAS gene sets, subtle polygenic effects, and regulatory SNPs in non-coding regions. These challenges often weaken enrichment signals and obscure direct gene-trait associations. To overcome these limitations, RNA-seq, proteomic, and expression data from previous studies were incorporated to validate candidate gene activity. This multi-tiered enrichment approach revealed a comprehensive molecular blueprint: KEGG pathways highlighted metabolic vigor and immune signaling (carbohydrate metabolism, Wnt, Toll, MAPK), while keyword and GO annotations emphasized signaling, innate immunity, stimulus response, organismal development, redox regulation, and chitin remodeling. Taken together, these findings illustrate a convergent adaptive strategy in Varroa-resistant phenotypes that combines metabolic reprogramming, enzymatic defense, and cellular signaling. This integrated framework underscores the complexity of honeybee resistance mechanisms and provides a foundation for functional validation studies and applied breeding strategies aimed at enhancing colony resilience.

These findings align strongly with the Omnigenic model, which posits that complex traits such as resistance to Varroa infestation are influenced not only by core genes with direct biological functions but also by peripheral genes that modulate trait expression through interconnected regulatory networks. The observed diversity of enriched pathways and molecular functions reflects widespread genomic involvement, where even genes with seemingly indirect roles contribute cumulatively to trait resilience via regulatory cascades and molecular cross-talk. This integrative view suggests that Varroa resistance arises from a network-wide genetic architecture, supporting the idea that broadly expressed and interconnected genes shape adaptive phenotypes in concert with core functional loci (Boyle et al. 2017; Mathieson 2021; Davoodi et al. 2022a; Davoodi & Razmkabir 2025).

## Conclusion

This study advances our understanding of the genetic architecture underlying hygienic behavior in honey bees, specifically the uncapping and removal of *Varroa*-infested brood cells. Genome-wide association analyses identified SNPs linked to this trait and enrichment analyses anchored these associations within broad functional networks spanning metabolic, immunological, and signaling pathways. The convergence of gene annotations supports the view that *Varroa* resistance is orchestrated by a distributed set of both core and peripheral genes, consistent with the Omnigenic model. This framework highlights the role of widespread regulatory interactions, extending beyond canonical effectors, and underscores the systems-level nature of uncapping behavior as a social immunity trait shaped by dense molecular interplay. By elucidating the genetic determinants of uncapping behavior, our findings provide a foundation for functional validation studies and evolutionary analyses of resistance mechanisms. Importantly, the identification of robust genomic markers offers practical tools for marker-assisted selection, enabling the development of mite-resistant honey bee lines. In the face of global pollinator declines and persistent environmental challenges, such integrative genetic approaches represent a critical step toward sustainable apiculture and the long-term resilience of honey bee populations.

## Declarations

### Data Availability Statement

The datasets used in this study are publicly available at Dryad repository with following link https://datadryad.org/dataset/doi:10.5061/dryad.8635cs4h.

## Acknowledgments

The authors gratefully acknowledge financial support provided by the Vice President for Research and Innovation at the University of Kurdistan under post-doctoral contract number 22600/9/3/ص. This support, totaling $1500 for one year, was essential in facilitating the research presented in this study.

Also, we disclose moderate use of AI-assisted language editing tools to paraphrase and enhance the clarity of the manuscript text. All scientific interpretations, analyses, and conclusions remain the responsibility of the authors.

## Authors’ contributions

All the authors contributed significantly to this study. PD contributed to data quality control, methodology development, data analysis, interpretation, and handled manuscript drafting and revision. MR provided guidance, and critical feedback throughout the research process. Both authors reviewed and approved the final version of the manuscript and agreed to be accountable for all aspects of the work.

## Ethics approval

This study did not require ethics approval, as it exclusively utilized publicly available data. No human participants, animals, or personally identifiable information was involved in the research. All the data were obtained from open-access sources and referenced completely.

## Consent for Publication

All the authors and participants involved in this study provided explicit consent for the publication of this research in G3 Journal.

